# Computational screening of T-muurolol for an alternative antibacterial solution against *Staphylococcus aureus* infections: A state-of-the-art phytochemical-based drug discovery approach

**DOI:** 10.1101/2024.06.07.597877

**Authors:** Soham Bhattacharya, Pijush Kanti Khanra, Adrish Dutta, Neha Gupta, Zahra Aliakbar Tehrani, Lucie Severová, Karel Šrédl, Marek Dvořák, Eloy Fernandez-Cusimamani

## Abstract

*Staphylococcus aureus* infections present a significant threat to the global healthcare system. The increasing resistance to existing antibiotics and their limited efficacy underscores the urgent need to identify new antibacterial agents with low toxicity to effectively combat various *S. aureus* infections. Hence, in this study, we have screened T-muurolol for possible interactions with several *S. aureus*-specific bacterial proteins to establish its potential as an alternative antibacterial agent. Based on binding affinity and interactions with amino acids T-muurolol was identified as a potential inhibitor of *S. aureus* lipase, dihydrofolate reductase, penicillin-binding protein 2a, D-Ala:D-Ala ligase, and RPP TetM, which indicates its potentiality against *S. aureus* and its multi-drug resistant strains. Also, T-muurolol exhibited good antioxidant and anti-inflammatory activity by showing strong binding interactions with FAD-dependent NAD(P)H oxidase, and cyclooxygenase-2. Consequently, MD simulation and recalculating binding free energies elucidated its binding interaction stability with targeted proteins. Furthermore, quantum chemical structure analysis based on density functional theory (DFT) depicted a higher E_HOMO-LUMO_ energy gap with a lower chemical potential index, and moderate electrophilicity suggests its chemical hardness and stability and less polarizability and reactivity. Additionally, pharmacological parameters based on ADMET, Lipinski’s rules, and bioactivity score validated it as a promising drug candidate with high activity toward ion channel modulators, nuclear receptor ligands, and enzyme inhibitors. In conclusion, the current findings suggest T-muurolol as a promising alternative antibacterial agent that might be a potential phytochemical-based drug against *S. aureus*. This study also suggests further clinical research before human application.

**Author Summary:** *Staphylococcus aureus* significantly contributes to human mortality, with over 1 million deaths annually accredited to its infections. At the same time, antimicrobial resistance (AMR) is a critical public health issue, responsible for an estimated 1.27 million deaths globally in 2019. The overuse and abuse of antimicrobials in both human and veterinary medicine are primary drivers of AMR, complicating the treatment of infections and increasing the risks associated with surgeries and other medical events. Despite the availability of antimicrobials such as methicillin, vancomycin, daptomycin, and linezolid, the emergence of multidrug-resistant *S. aureus* poses a formidable challenge to effective treatment. Due to the limited efficacy and increasing resilience to current antibiotics, there is an urgent need to discover new and effective antibacterial drugs against *S. aureus*. Since time immemorial, phytochemicals have been valued for their rich biological properties and safety in treating bacterial infections. In this study, we have computationally investigated T-muurolol as a potential alternative antibacterial agent. Our molecular docking and simulation approaches provide insights into the interactions of T-muurolol as an inhibitor of *S. aureus*-specific bacterial proteins. Additionally, pharmacokinetic and quantum chemical structure analyses offer valuable information about T-muurolol’s potential as a drug candidate, supporting its further development as an antibacterial agent.

## Introduction

The population growth and invasion dynamics of *S. aureus* are undoubtedly opportunistic as they subvert the human immune surveillance machinery by evoking the metastatic colony dissemination from superficial to life-threatening levels [1]. The source of bacterial dissemination mainly evolved from the mucosal contacts as the mucosal layer mainly favours harbouring the bacterial strains. Therefore, the mucosal surfaces of the nose, throat, vaginal wall, open wound, and gastrointestinal tract impart favourable colonization carriages of *S. aureus* bacteraemia (SAB) [2]. Colonization is also frequent in human juveniles and patients who have already been vulnerable to HIV and diabetes [3]. Despite the backdrops of prevalence and prognostic outcomes of SAB in the last decade, it has been witnessed the emergence of multi-drug resistant (MDR) strains to empiric therapies, antibiotics, and large-scale treatment options [4]. Although, the present array of antibiotics, including methicillin, vancomycin, daptomycin, and linezolid, multidrug-resistant *S. aureus* has emerged as the most formidable bacterial strain, causing life-threatening infections and illnesses in humans [5]. The estimated numbers of invasive disease cases and morbidity-mortality in the United States due to methicillin-resistant *S. aureus* (MRSA) pathogenesis were 94,360 and 18,650 in 2005, respectively [2]. Similarly, MRSA infection is associated with annual healthcare costs of about $3 billion a year and is set to increase over time [6]. The rampant and escalating severity of diseases, coupled with the failure of rational therapeutic approaches, due to the evolution of MDR *S. aureus* strains, might lead one to speculate that humans could become highly vulnerable and unmatched in facing the uncontrolled infectious threat posed by *S. aureus* [3]. Therefore, considering the systemic toxicity and limited efficacy of current antibacterial medications, identifying the underlying mechanisms of this epidemic and developing alternative antibacterial drugs should be the prime goal of the current research. These new drugs should aim to achieve high success rates against *S. aureus* and prevent obsolescence, thereby mitigating the uncertainty of public health and improving patient care for SAB-associated infections.

Plant-derived phytochemicals can be adopted as sustainable alternative antibacterial agents. Over the last three decades, phytochemicals have undergone extensive experimental validation, establishing themselves as emerging antibacterial agents with significant potential [7]. In the treatment of penicillin G-resistant strains of *S. aureus*, many phytochemicals have successfully passed experimental tests against various MDR strains [8]. The activity of phytochemicals-based antibacterial not only possesses antibacterial activity but also acts as an anti-inflammatory, and antioxidant agent that can dramatically improve the targeting of the infection site while minimizing systemic exposure and associated toxicity [9]. Therefore, different classes of phytochemicals such as phenols, alkaloids, terpenoids, flavonoids, carotenoids, organosulfur compounds, and coumarins, etc. are actively available in the market as FDA-approved drugs and used in herbal medicinal products [10]. Phytochemicals have been shown to impede major bacterial MDR factors, including bacterial virulence factors, ion pumps, bacterial cell wall toxins, cellular microstructure, replication machinery, and membrane permeability [11]. Therefore, this study focuses on the *in-silico* repurposing of a new generation of phytochemicals against *S. aureus*, aiming to explore alternative antibacterial therapeutic approaches.

T-muurolol is a cadinane sesquiterpenoid with versatile characteristics, as several studies have demonstrated its diverse biological activities, including antimicrobial, antioxidant, anti-inflammatory, and antitermitic effects [12, 13, 14]. Although T-muurolol is a potential candidate with various reported antimicrobial activities and could serve as an alternative solution to combat *S. aureus* infections, the phytochemical-based drug-reprofiling approach is still in its early stages. Currently, there are no such research articles detailing the re-profiling of T-muurolol as an alternative treatment for suppressing SAB-associated infections, and also, its molecular interaction with *S. aureus*-associated proteins remains largely unexplored. Therefore, this study investigates the potential of T-muurolol as an anti-staphylococcal agent through molecular docking, pharmacokinetics, molecular dynamics, and chemical structure analysis.

## Results

### Molecular docking and interaction analysis

The molecular interactions between the T-muurolol and tested bacteria-specific proteins for *S. aureus* along with various proteins related to antioxidant activity and inflammation in humans are summarized in Table 1. T-muurolol exhibited a good binding affinity with all the proteins used in the study between the range of -7.5 kcal/mol to -4.3 kcal/mol. T-muurolol demonstrated a strong binding affinity for *S. aureus* lipase (6KSI), showing a binding energy of -7.1 kcal/mol, which is notably higher than its affinity for other tested pathogenic proteins. A hydrogen bond of bond length 3.78 Ǻ with A chains of amino acid residue Asp236 along with non-weak interactions with A chains of Leu250, Lys249, and Tyr240 was found in this interaction (Fig 1C and 1D). T-muurolol also demonstrated significant binding interactions (with a binding energy of -7.5 kcal/mol) with dihydrofolate reductase (3SRW), a key enzyme responsible for bacterial DNA synthesis. Notably, T-muurolol formed a hydrogen bond of 5.03 Å with amino acid Phe93, along with other interactions involving Ala8, Leu6, Leu21, Phe93, and Val32 (Fig 1A and 1B). Similarly, for MDR *S. aureus* penicillin-binding protein 2a (1MWU) is a key drug target responsible for broad-spectrum beta-lactam resistance. T-muurolol also showed a strong binding interaction with 1MWU (-6.1 kcal/mol) with a hydrogen bond of 4.42 Ǻ with amino acid residue Glu294 and other interactions with Ala276, Lys289, and Tyr272 (Fig 2A and 2B). The robust interactions between PBPs and T-muurolol can effectively inhibit MRSA by disrupting peptidoglycan cross-linking and impeding bacterial cell wall synthesis. Likewise, D-Ala:D-Ala ligase (3N8D), a potential target for VRSA due to its role in ligase impairment, is affected by T-muurolol, evidenced by significant binding energy with 3N8D (-5.9 kcal/mol). Furthermore, T-muurolol demonstrates substantial binding affinities with RPP TetM (3J25) with a binding energy of -6.6 kcal/mol. These proteins serve as key targets in the context of TetRSA, emphasizing T-muurolol’s potential as a therapeutic agent against resistant bacterial strains.

**Table 1.**
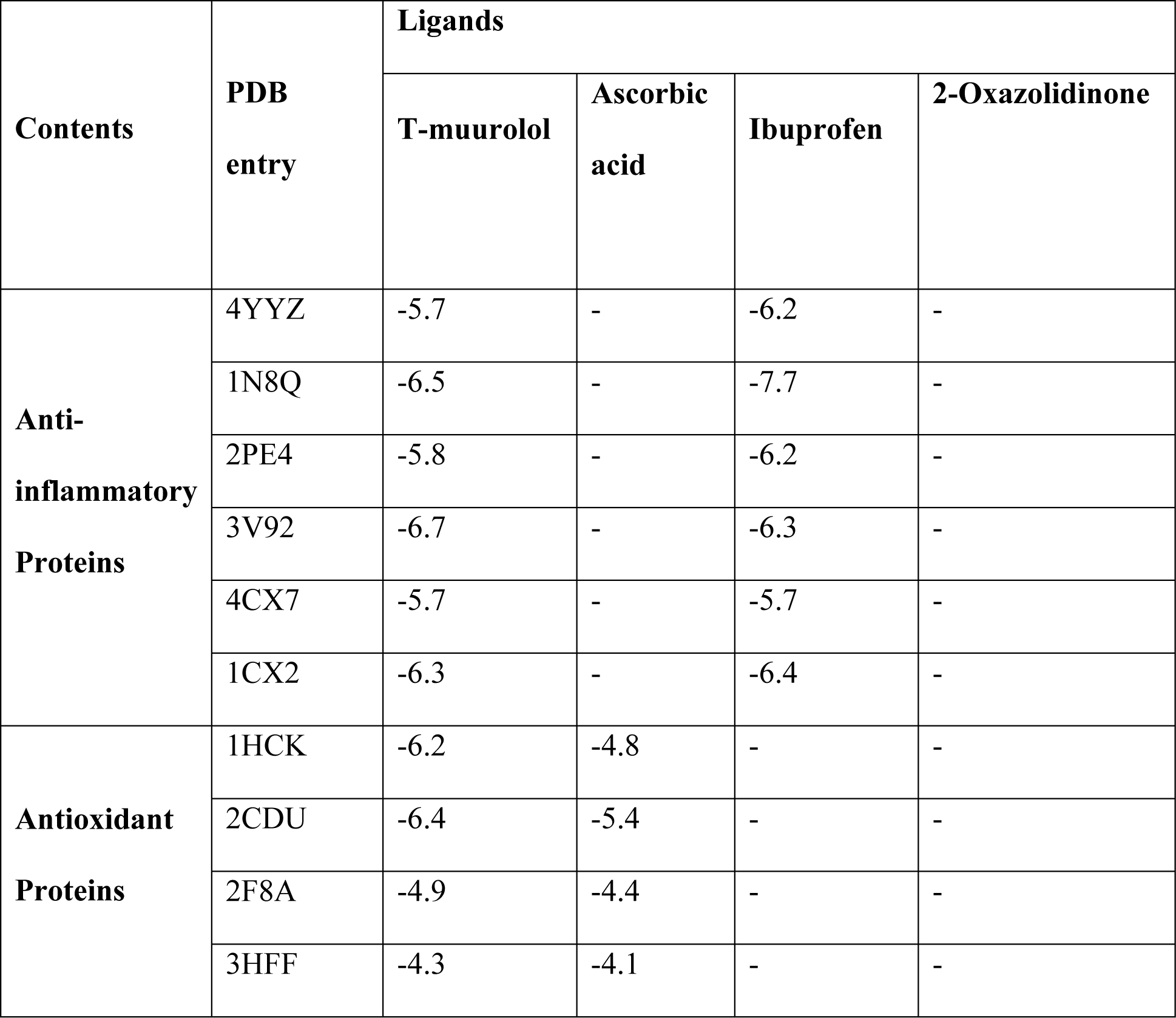
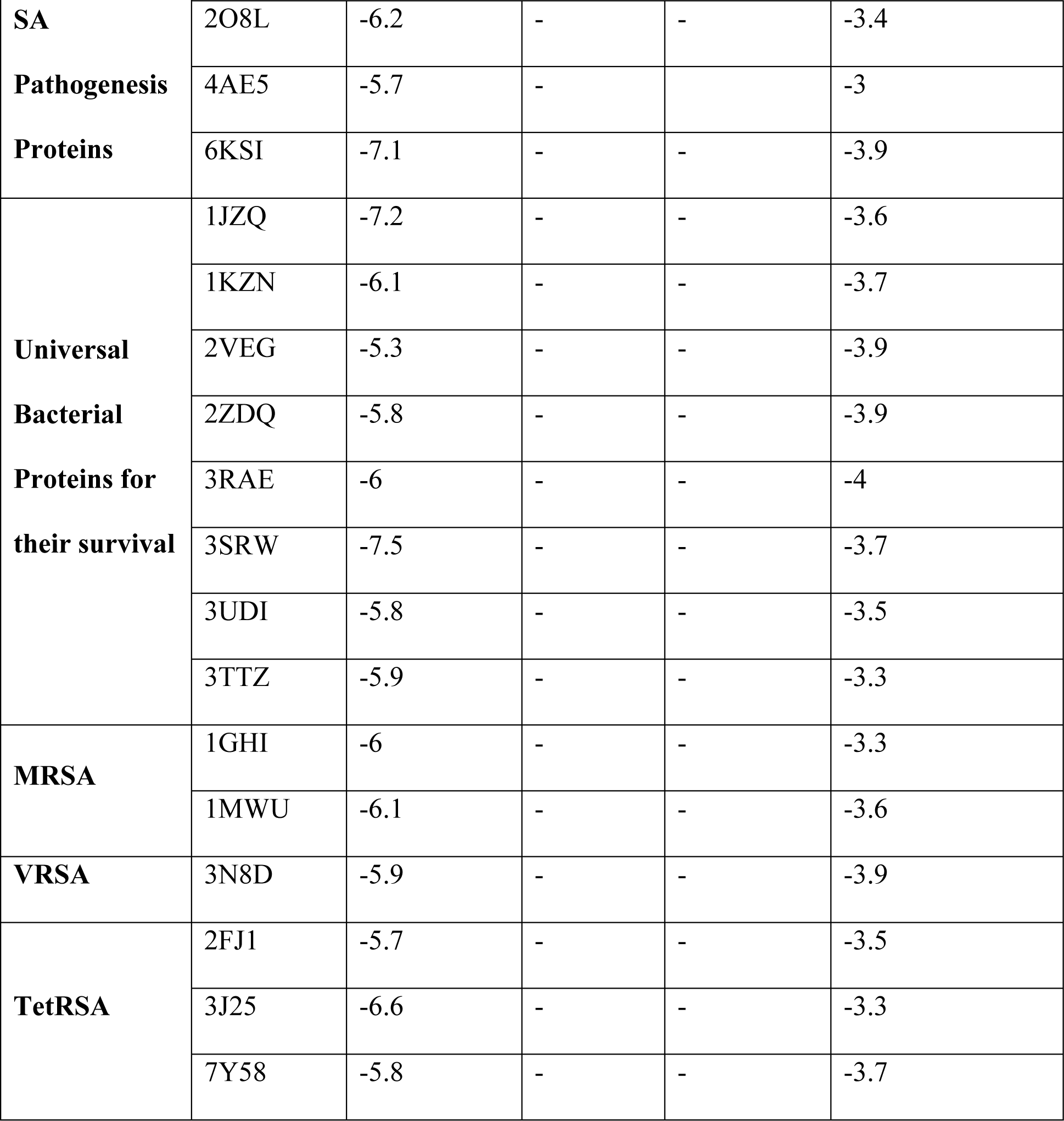
Binding free-energy values (kcal/mol) of T-muurolol as ligand along with ascorbic acid, ibuprofen and 2-oxazolidinone as references for good antioxidant, anti-inflammatory and antibacterial agents, respectively.

**Table 2.** Top hit interactions of T-muurolol with proteins and their hydrogen bonds and several other interactions.

| **PDB**<br>**entry** | **No. of**<br>**H-**<br>**bonds** | **H-bonds and**<br>**interacting**<br>**residues** | **No. of other**<br>**interactions** | **Other**<br>**interactions and**<br>**numbers** | **Other**<br>**interaction**<br>**and**<br>**interacting** |
| --- | --- | --- | --- | --- | --- |

|  |  |  |  |  | **residues** |
| --- | --- | --- | --- | --- | --- |
| 1CX2 | 0 |  | 8 | Alkyl and Pi-alkyl (8) | Ile112 (2),<br>Tyr115 (2),<br>Val89 (3),<br>Val116 (1) |
| 1MWU | 1 | Glu294 (1) | 6 | Alkyl and Pi-alkyl (5), Pi-sigma (1) | Ala276 (1),<br>Lys289 (3),<br>Tyr272 (2) |
| 2CDU | 0 |  | 11 | Alkyl and Pi-alkyl (10), Pi-sigma (1) | Ile160 (2),<br>Leu299 (2),<br>Pro298 (1),<br>Tyr159 (3),<br>Tyr188 (3) |
| 3J25 | 0 |  | 10 | Alkyl and Pi-alkyl (10) | Ala554 (2),<br>Cys603 (2),<br>Ile544 (2),<br>Leu593 (4) |
| 3N8D | 0 |  | 2 | Alkyl (2) | Val19 (2) |
| 3SRW | 1 | Phe93 (1) | 11 | Alkyl and Pi-alkyl (10), Pi-sigma (1) | Ala8 (2),<br>Leu6 (2),<br>Leu21 (3),<br>Phe93 (2),<br>Val32 (2) |
| 6KSI | 1 | Asp236 (1) | 6 | Alkyl and Pi-alkyl (6) | Leu250 (1),<br>Lys249 (3), |
|  |  |  |  |  | Tyr240 (2) |

**Fig 1.**
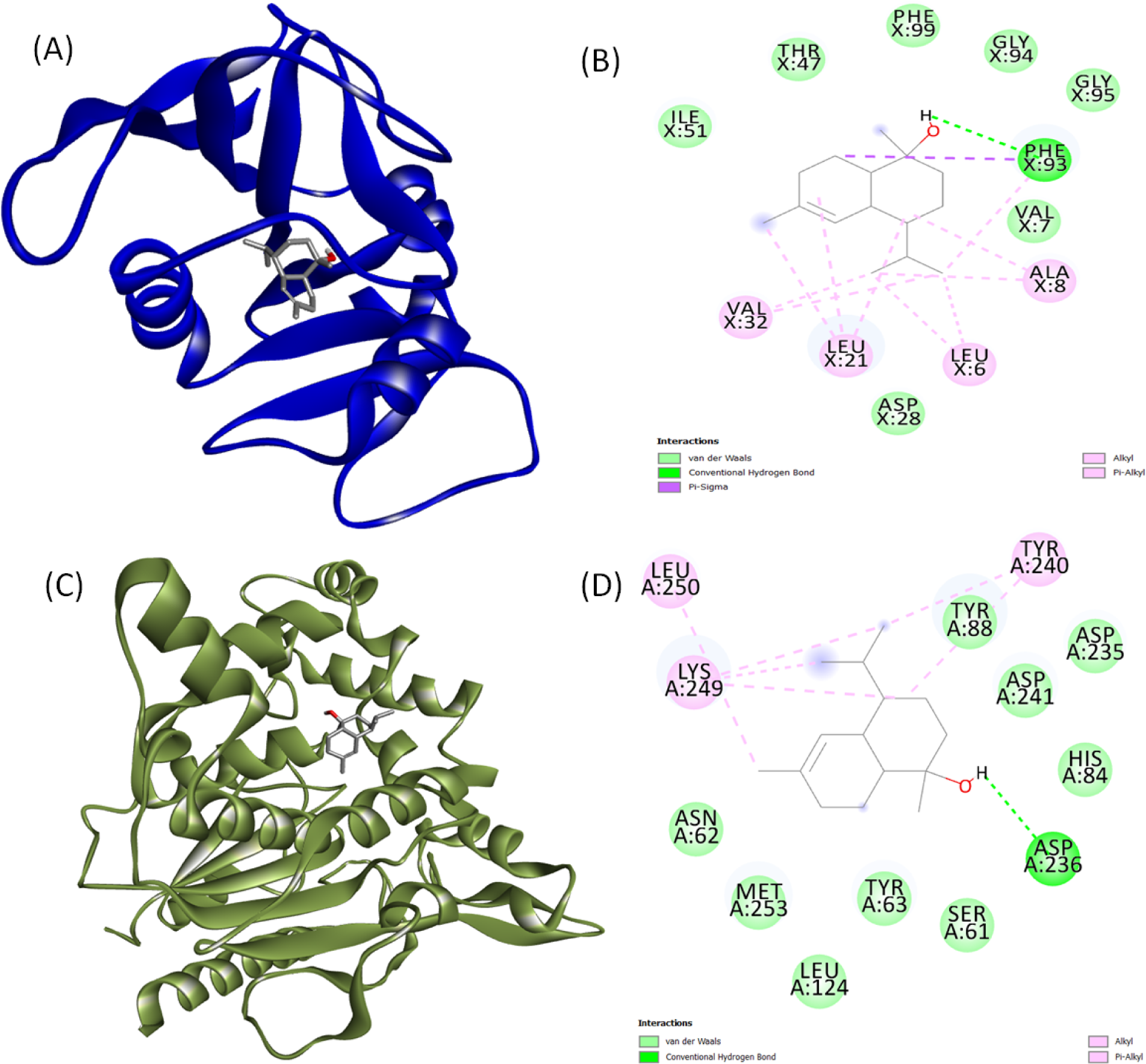
3D and 2D interactions of T-muurolol as ligand with (A) and (B) Dihydrofolate reductase (PDB ID:3SRW), and (C) and (D) S. aureus lipase (PDB ID: 6KSI) as proteins.

**Fig 2.**
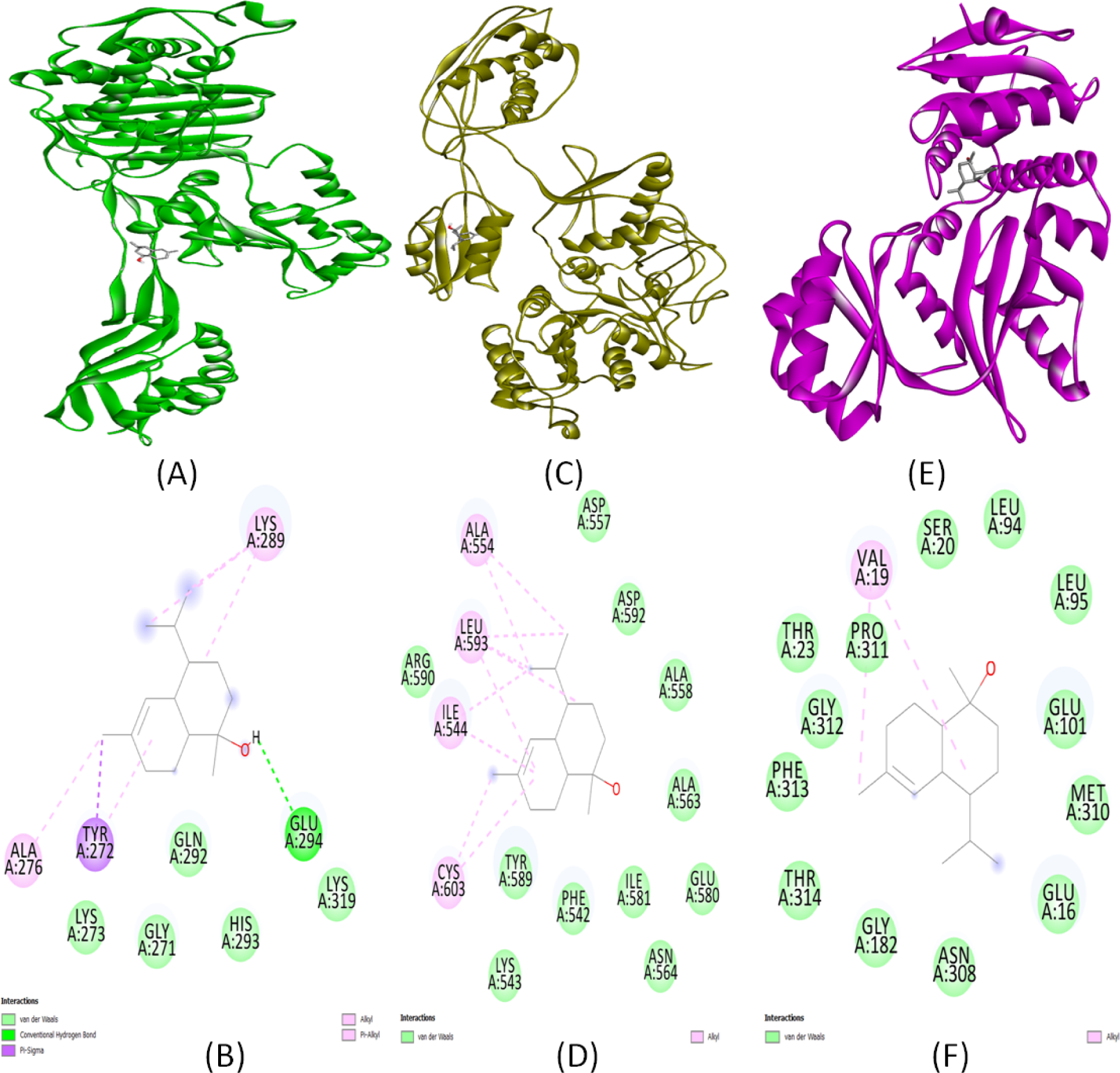
3D and 2D interactions of T-muurolol as ligand with (A) and (B) penicillin-binding protein 2a (PDB ID: 1MWU), (C) and (D) RPP TetM in complex with the 70S ribosome (PDB ID: 3J25), and (E) and (F) D-Ala:D-Ala ligase (PDB ID: 3N8D) as proteins.

**Fig 3.**
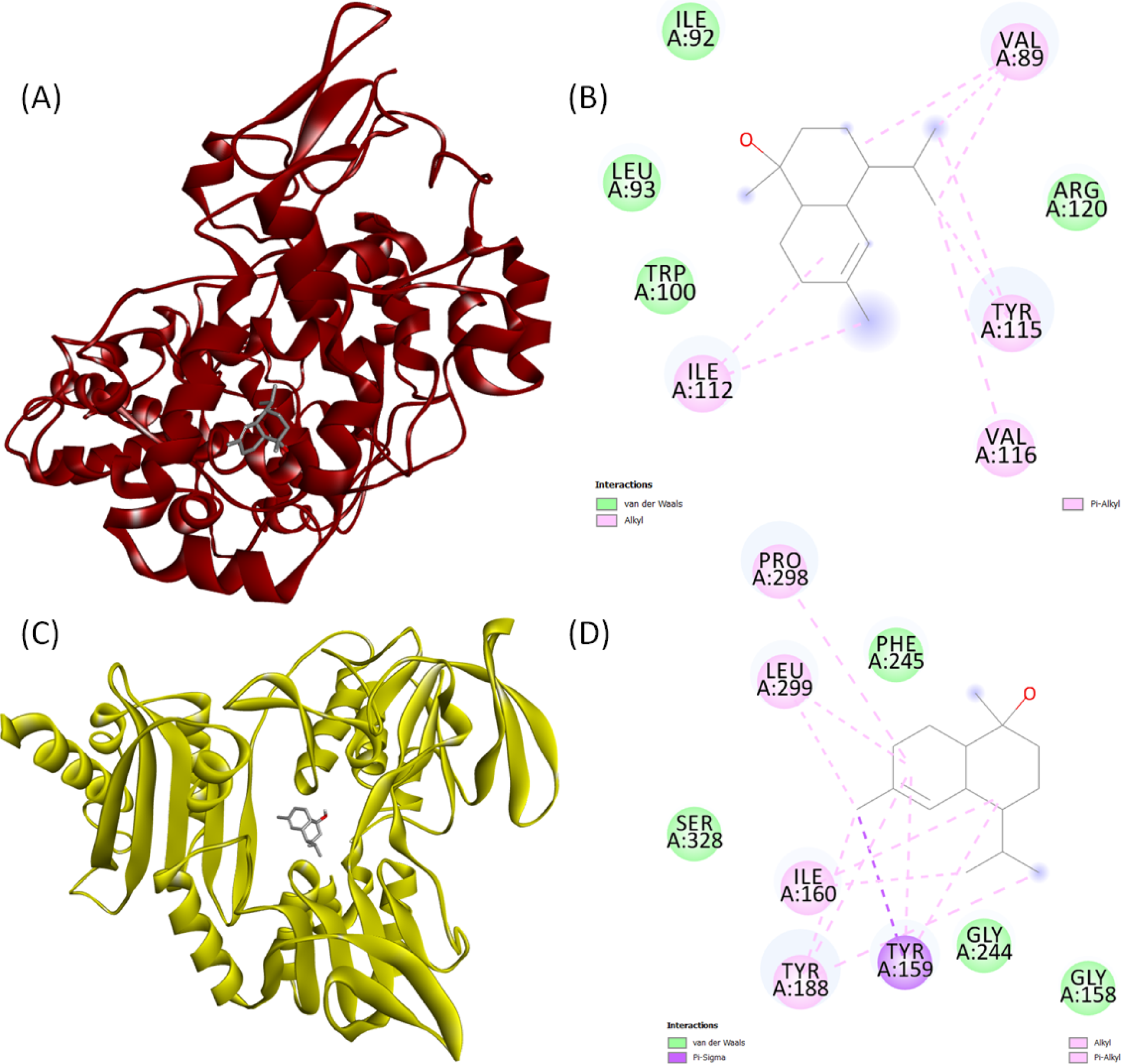
**3D and 2D interactions of T-muurolol as ligand with (A) and (B) cyclooxygenase-2 (PDB ID: 1CX2), and (C) and (D) FAD-dependent NAD(P)H oxidase (PDB ID: 2CDU) as proteins.**

Moreover, T-muurolol displayed favourable binding interactions with FAD-dependent NAD(P)H oxidase(2CDU) (-6.4 kcal/mol) among all the tested antioxidant proteins. Additionally, it demonstrated strong binding affinity with cyclooxygenase-2 (1CX2) (-6.3 kcal/mol) which was higher compared to the reference anti-inflammatory drug ibuprofen (Table 1).

### ADMET pharmacokinetic analysis

#### Absorption

The water solubility of tested compounds, cell permeability utilizing the colon carcinoma (Caco-2) cell line, human intestinal absorption, skin permeability, and whether the molecule is a P-glycoprotein substrate or inhibitor are the primary parameters for evaluating drug absorption criteria. Current results suggested that T-muurolol has lower solubility to water whereas the reference antibiotic drug 2-oxazolidinone seems to be highly soluble (Table 3). Caco-2 cell permeability influences the ultimate bioavailability, according to Chandra et al., (2021)[19], a drug with a value > 0.90 is deemed highly permeable. The obtained results exhibited the high permeability of T-muurolol in the Caco-2 cell line with a value of 1.479. Similarly, all the tested compounds showed a higher gastrointestinal (GI) absorption percentage which was more than 90% (Table 3). A study by Saha et al. (2021)[20] explained that the human intestine is primarily an active location for drug absorption, with more than 30% being considered rapidly absorbed. Our results also indicated that T-muurolol along with reference drugs is neither a P-glycoprotein substrate nor an inhibitor (Table 3).

**Table 3.**
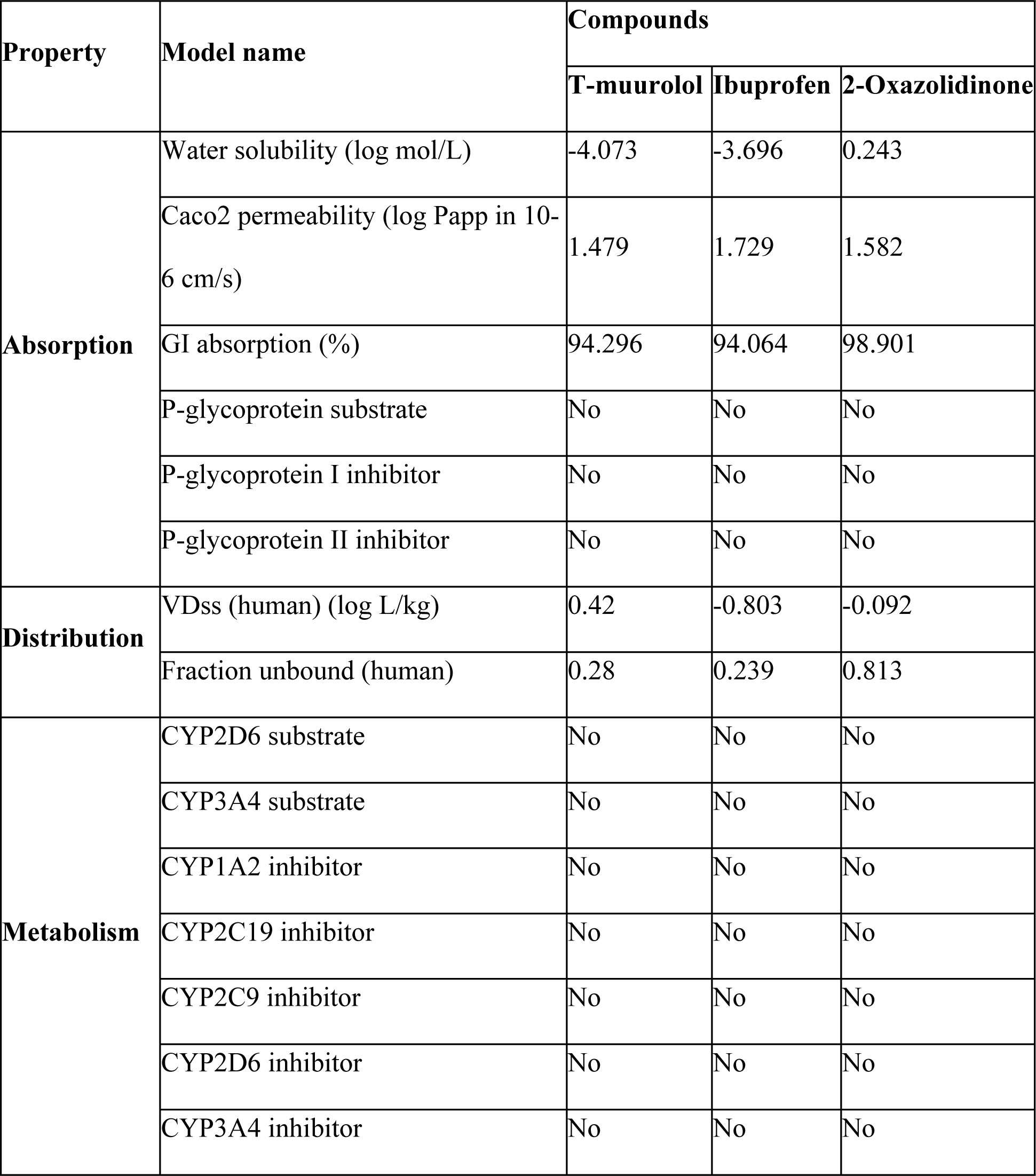
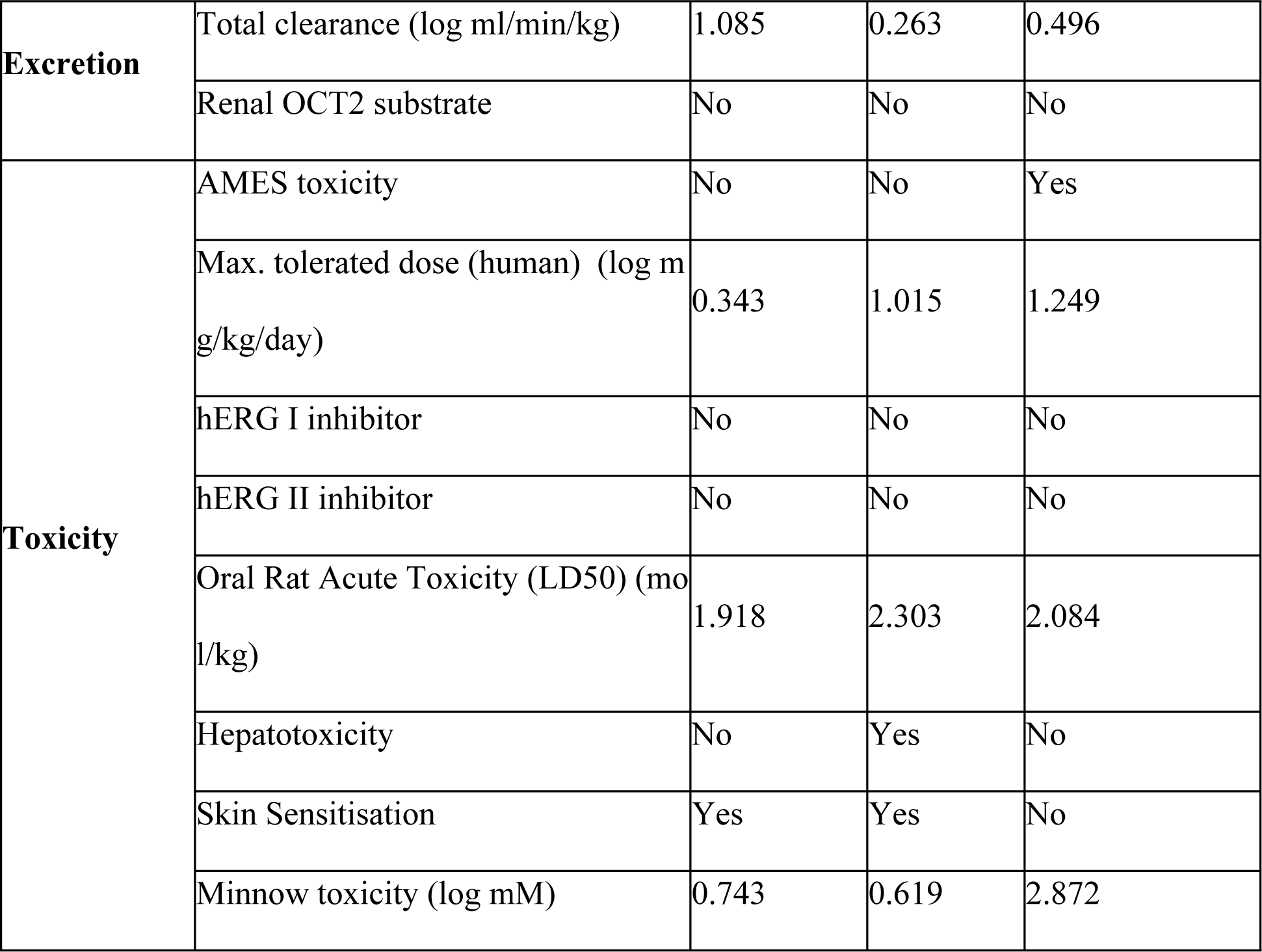
Predicted ADMET pharmacokinetic properties.

#### Distribution

Here drug distribution is generally analyzed based on the steady-state volume of distribution (VDss), and fraction of unbound value to human plasma. The steady-state volume of distribution (VDss) is a key pharmacokinetic parameter for determining drug dosages, indicating the theoretical volume needed to achieve the same blood plasma concentration as the administered dose. Higher VDss values suggest greater distribution into tissues rather than plasma. For antibiotics and antivirals, extensive tissue distribution is often desired. VDss is considered low if log(VDss) < -0.15 and high if >0.45 [20]. T-muurolol showed a moderate VDss value of 0.42 log L/kg which suggests its great distribution over tissue (Table 3). A drug’s effectiveness is influenced by its binding to blood proteins, more binding facilitates membrane crossing [21]. The fraction unbound to human plasma should be between 0.02 and 1.0 [21]. T-muurolol showed good fraction unbound values with 0.28 (Table 3).

#### Metabolism

The Cytochrome P450 (CYP) enzyme group includes isozymes that metabolize drugs, fatty acids, steroids, bile acids, and carcinogens. Drug metabolism depends on whether a substance is a CYP substrate or inhibitor. It was study found that T-muurolol and the reference drugs tested are neither substrates nor inhibitors of CYP enzymes (Table 3), indicating they will be metabolized effectively and not hindered by the body’s biological processes.

#### Excretion

Organic cation transporter 2 (OCT2) is a renal uptake transporter crucial for drug deposition and clearance from the kidneys [19]. Excretion is assessed by total clearance and whether the compound is a substrate of OCT2 or not. The predicted results showed that T-muurolol is not a substrate of OCT2 (Table 3), suggesting it may be eliminated through a different route. Also, T-muurolol showed a moderate excretion clearance efficacy with a log(CLtot) value of 1.085 ml/min/kg.

#### Toxicity

The toxicity of the drug compounds was assessed using AMES results, human maximum tolerated dose, oral rat-acute toxicity, hepatotoxicity, skin sensitization, minnow toxicity, and human ether-a-go-go gene (hERG) inhibition. The AMES result indicated that T-muurolol is non-mutagenic and non-carcinogenic (Table 3). The Maximum Recommended Tolerance Dose (MRTD) estimates human toxic doses, considered low if below log 0.477 (mg/kg/day) [20]. So, T-muurolol showed less toxicity with a MRTD value of log 0.343 mg/kg/day whereas reference drugs seem to be highly toxic. The results also showed that none of the tested compounds are non-hERG inhibitors, however, T-muurolol was predicted to be skin sensitive although no hepatotoxic prediction was established. All of the tested compounds showed higher oral rat-acute toxicity (LD_50_) (Table 3) which indicated less lethality compared to one with a lower LD_50_ value [22]. If a molecule’s log LC50 (concentration causing 50% fathead minnow mortality) is below 0.5 mM (log LC_50_ < -0.3), it’s deemed highly acutely toxic. Current findings suggest that T-muurolol and reference drugs are less toxic, with significantly higher scores than the mentioned LC_50_ threshold (table 3).

### Drug-likeness and bio-activity analysis

The drug-likeness property of the compounds is based on Lipinski’s rule of five which encompasses crucial molecular properties that impact a drug’s pharmacokinetics in the human body. Current results showed that none of the tested compounds violated Lipinski parameters such as molecular weight, number of hydrogen bond donors and acceptors, and octanol-water partition coefficient (Table 4). Topological polar surface area (TPSA) calculation indicates a drug’s bioavailability and its hydrogen bonding potential. All tested compounds have a TPSA range of 20.23–38.33 Å (Table 4), significantly lower than the typical range of 160 Å [23]. Our results also indicated an ideal bioavailability score of 0.55 for T-muurolol which will be well-absorbed by the human body [24].

**Table 4.** Drug likeness properties.

| Compounds | Lipinski | Molecular Weight (g/mol) | H-bond acceptors | H-bond donors | LogP | TPSA (Å<sup>2</sup>) | Bioavailability score |
| --- | --- | --- | --- | --- | --- | --- | --- |
| T-muurolol | Yes; 0 violation | 222.37 | 1 | 1 | 3.42 | 20.23 | 0.55 |
| Ibuprofen | Yes; 0 violation | 206.28 | 2 | 1 | 3.01 | 37.3 | 0.85 |
| 2-Oxazolidinone | Yes; 0 violation | 87.08 | 2 | 1 | -0.04 | 38.33 | 0.55 |

Bioactivity scores for the drug compound can be calculated based on parameters including kinase inhibition, protease inhibition, enzyme activity inhibition, binding to G protein-coupled receptors (GPCRs) and nuclear receptors, and ion channel modulation. The bioactivity score indicated that T-muurolol is highly active toward ion channel modulators, nuclear receptor ligands, and enzyme inhibitors with moderate activity toward GPCR ligands, but at the same time, it is inactive toward kinase inhibitors and protease inhibitors (Table 5).

**Table 5.** Prediction of bioactivity score of compounds.

| Compounds | *GPCRs | ICM | KI | NRL | PI | EI |
| --- | --- | --- | --- | --- | --- | --- |
| T-muurolol | -0.09 | 0.05 | -0.87 | 0.39 | -0.63 | 0.4 |
| Ibuprofen | -0.17 | -0.01 | -0.72 | 0.05 | -0.21 | 0.12 |
| 2-Oxazolidinone | -3.47 | -3.26 | -3.67 | -3.78 | -3.23 | -3.4 |

\*GPCRs G protein-coupled receptors, ICM ion channel modulator, KI kinase inhibitor, NRL nuclear receptor ligand, PI protease inhibitor, EI enzyme inhibitor. A molecule with a bioactivity score above 0.00 is likely to exhibit significant biological activities; scores ranging from -0.50 to 0.00 are considered moderately active, while those below -0.50 are presumed inactive.

### MD simulation analysis

Structural integrity, stability, compactness protein folding of apo-protein (ligand-free protein) and holo-proteins (proteins with ligands) had been critically analysed based on some variable characteristics RMSD (root mean square deviation), RMSF (root mean square fluctuation), R_g_ (radius of gyration), Hydrogen bonds (HB) formation, SASA (solvent accessible surface area) and binding free energy decomposition (MM-PBSA) after 100 ns of MD run. The superimposition of T-muurolol on the targeted proteins was shown in Fig 4 to observe the time-dependent protein-drug interaction.

**Fig 4.**
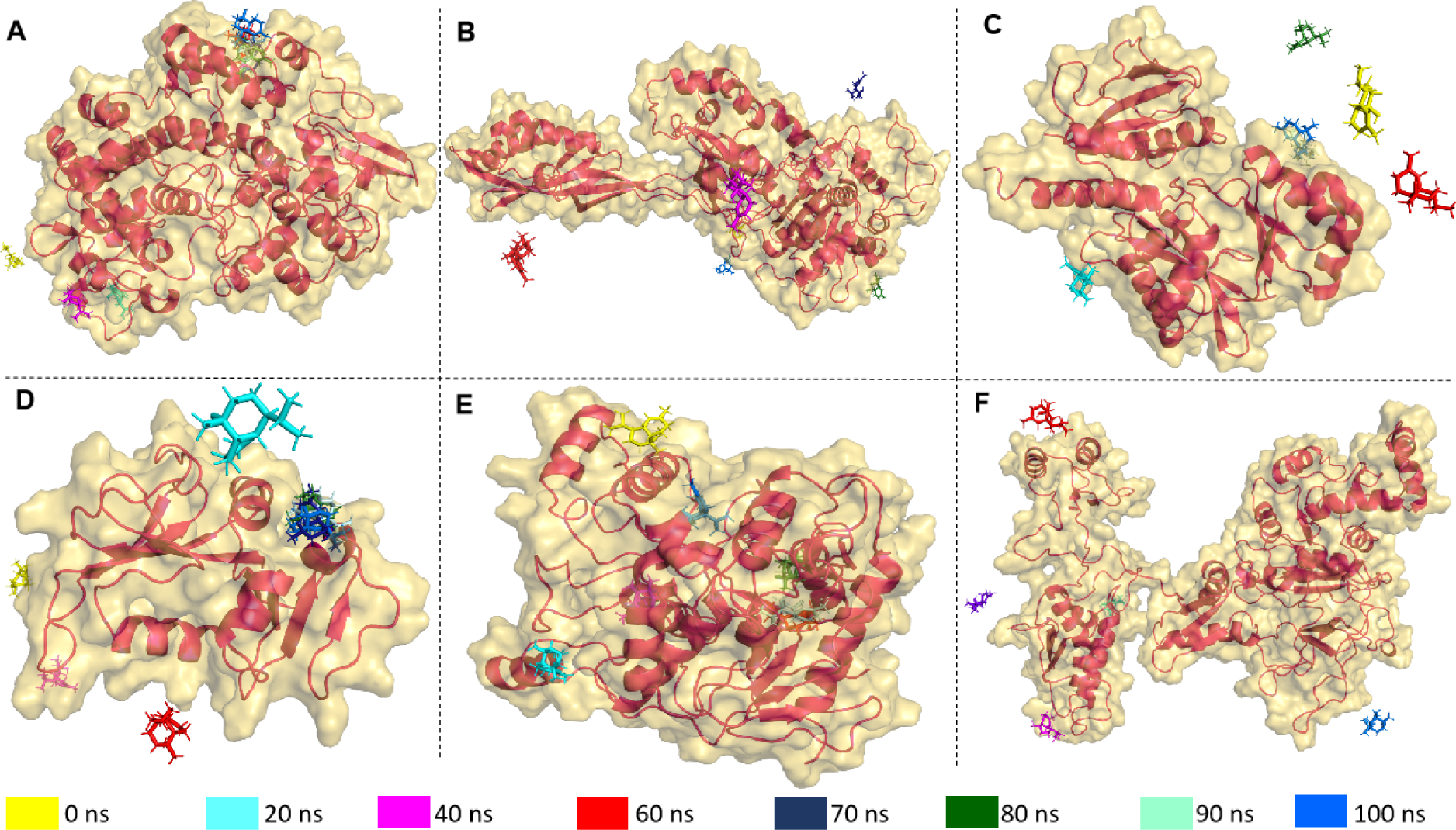
Superimposition of the selected drug T-muurolol on the targeted proteins (A) 1CX2 (B) 1MWU (C) 3N8D (D) 3SRW (E) 6KSI (F) 3J25 in the time-dependent manner of 10/ 20 ns intervals to observe the concomitant protein-drug interactions.

#### Monitoring conformational stability of the backbone

Root mean square deviation (RMSD) of the 6 apo-protein and holo-protein system was illustrated in Fig 5 where the apo-protein and holo-protein system of 1CX2, 1MWU, 3N8D, 3SRW, 6KSI and 3J25 range in between ∼(0.15-0.45) nm (subfigure A) ∼(0.18-0.64) nm (subfigure B), ∼(0.10-0.50) nm (subfigure C), ∼(0.05-0.27) nm (subfigure D), ∼(0.10-0.30) nm (subfigure E) and ∼(0.50-2.0) nm (subfigure F) with an average apo-proteins’ RMSD of ∼0.27 nm (1CX2), ∼0.34 nm (1MWU), ∼0.27 nm (3N8D), ∼0.15 nm (3SRW), ∼0.18 nm (6KSI) and ∼0.93 nm (3J25). In MD simulations, RMSD indicates the conformational stability of the C_α_ backbone and protein-ligand interactions. As shown in Fig 5A, holo-protein systems 1CX2-ibuprofen and 1CX2-T-muurolol deviated after 10 ns. The RMSD for 1CX2-ibuprofen was more erratic, suggesting greater backbone stability for 1CX2-T-muurolol throughout the MD trajectory. Similarly, in Figs 5E and 5F, aberrant fluctuations were observed for 6KSI and 3J25 holo-proteins between 0-60 ns and 10-100 ns, respectively, compared to their apo-proteins. This indicates that significant ligand-protein interactions can cause instability and deviations, leading to decreased conformational backbone stability. Conversely, Figs 5B and 5D show that the RMSD trajectories of 1MWU and 3SRW holo-proteins aligned closely with their apo-proteins, indicating that these proteins maintained the backbone stability upon ligand interaction. In Fig 5C, the comparative RMSD analysis reveals that the dip in RMS deviation of 3N8D-T-muurolol beyond that of 3N8D after 40 ns suggests π–π stacking between T-muurolol and 3N8D [25].

**Fig 5.**
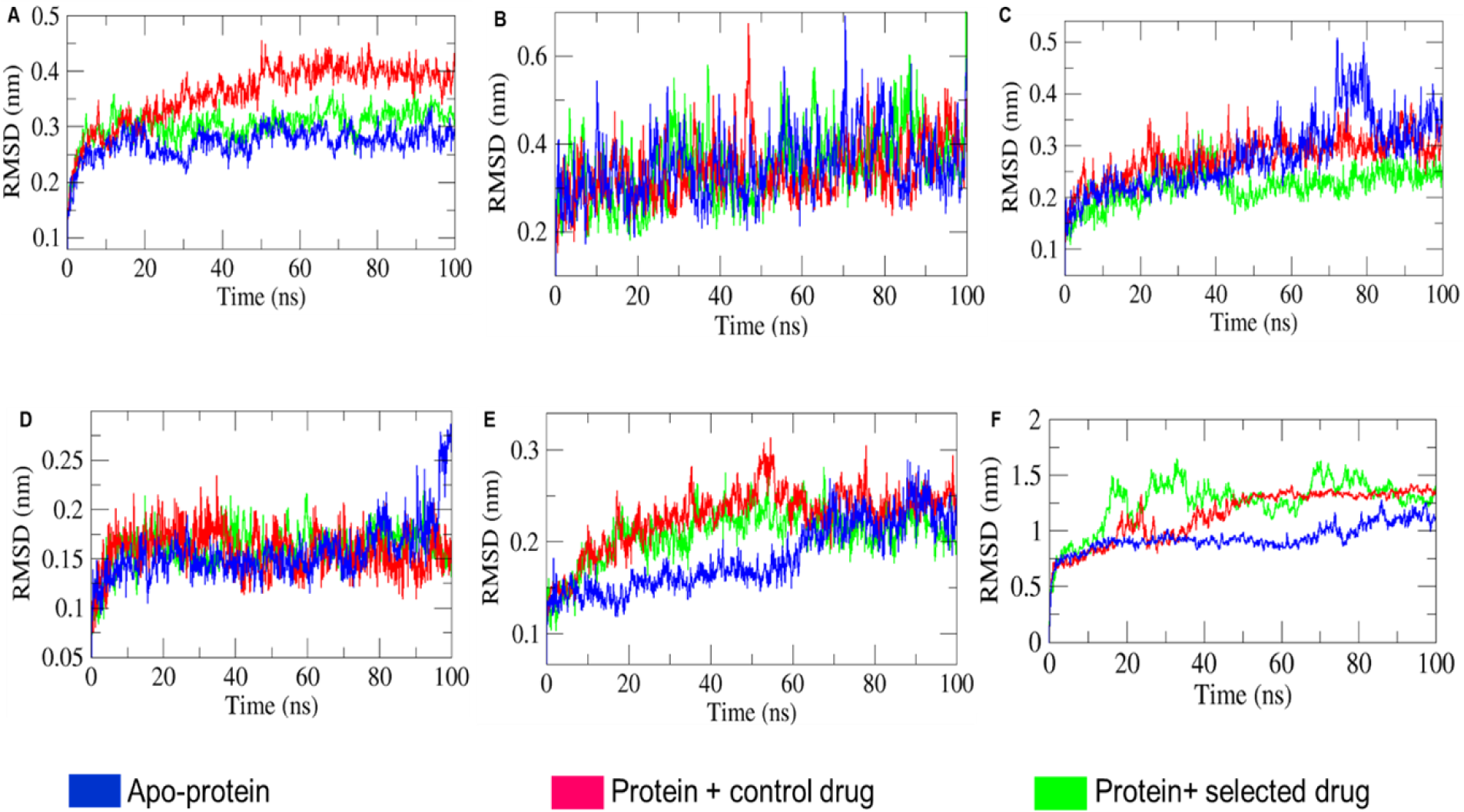
Comparative analysis of root-mean-square deviation (RMSD) of the consecutive apo-proteins and holo-proteins combinations (proteins with control drug and selected drug) mentioned in different subfigures (A) 1CX2 (B) 1MWU (C) 3N8D (D) 3SRW (E) 6KSI and (F) 3J25 combinations.

#### Residual flexibility analysis

Thermodynamic stability of protein-ligand complexes is determined by residual fluctuations (RMSF) of proteins in the presence of ligands. Comparing the RMSF of apo-proteins and holo-proteins characterizes the conformational stability and flexibility of amino acid Cα atoms. Larger RMSF peaks in holo-protein residues indicate greater flexibility, while smaller peaks indicate rigidity, reflecting maximal and minimal interactions with their ligands and surrounding polar molecules, respectively [25]. Fig 6 gives us the rationale insights about the comparative analysis of apo-protein and holo-protein specific average RMSF values which ranges as 0.17 ± 0.02 nm for 1CX2, 0.26 ± 0.001 nm for 1MWU, 0.21 ± 0.04 nm for 3N8D, 0.13 ± 0.01 nm for 3SRW, 0.13 ± 0.005 nm for 6KSI and 0.32 ± 0.24 nm for 3J25. From the average RMSF values of combined apo-protein and holo-protein systems, 3J25 holo-protein showed the largest increase in fluctuation, while 6KSI holo-protein showed the smallest change compared to their apo-proteins. This indicates that 3J25 maintained residual flexibility, whereas 6KSI retained rigidity in the presence of specific ligands. Among the six holo-proteins, 6KSI demonstrated the best interactions with ligands and polar molecules, while 3J25 showed the least interaction, as supported by binding free energy calculations in Table 6 for T-muurolol and protein interactions. Fig 6 shows that higher RMSF values for specific holo-protein residues 80-100 for 1CX2-ibuprofen, 390-410 for 1MWU-T-muurolol, 75-78 for 3N8D-2-Oxazolidinone, 175-240 for 6KSI-2-Oxazolidinone, 400-550 for 3J25-2-Oxazolidinone, and 550-620 for 3J25-T-muurolol indicate that proteins associated with T-muurolol maintained greater residual rigidity compared to their control drug complexes. This suggests that T-muurolol exhibited more stable interactions with all proteins over the 100 ns MD timespan compared to the control drugs.

**Table 6.** Binding free energy (KJ/mol) calculation of the following proteins based on MD interaction with T-muurolol.

| Macromolecules | Van der<br>waals energy<br>(KJ/mol) | Electrostatic<br>energy<br>(KJ/mol) | Polar<br>solvation<br>energy<br>(KJ/mol) | SASA<br>energy<br>(KJ/mol) | Binding<br>free<br>energy<br>(KJ/mol) |
| --- | --- | --- | --- | --- | --- |
| 1CX2 | $-53.91 \pm 34.59$ | $-2.68 \pm 3.76$ | $23.93 \pm 30.61$ | $-6.82 \pm 5.92$ | $-39.48 \pm 49.61$ |
| 1MWU | $-0.05 \pm 0.12$ | $0.46 \pm 1.43$ | $-55.05 \pm 74.16$ | $-0.93 \pm 1.70$ | $-55.58 \pm 73.88$ |
| 3N8D | $-12.92 \pm$ | $0.58 \pm 2.59$ | $-6.58 \pm$ | $-2.46 \pm 4.00$ | $-21.39 \pm$ |
|  | 26.82 |  | 82.83 |  | 80.83 |
| 3SRW | -29.99 ±<br>38.37 | -1.68 ± 5.70 | 24.85 ±<br>38.82 | -4.21 ± 5.34 | -11.03 ±<br>50.33 |
| 6KSI | -71.82 ±<br>36.20 | -4.36 ± 7.12 | 36.27 ±<br>19.84 | -10.72 ±<br>5.37 | -50.64 ±<br>52.25 |
| 3J25 | -5.164 ± 12.72 | -0.14 ± 0.93 | 2.14 ± 64.56 | -1.84 ± 2.37 | -5.01 ±<br>67.25 |

**Fig 6.**
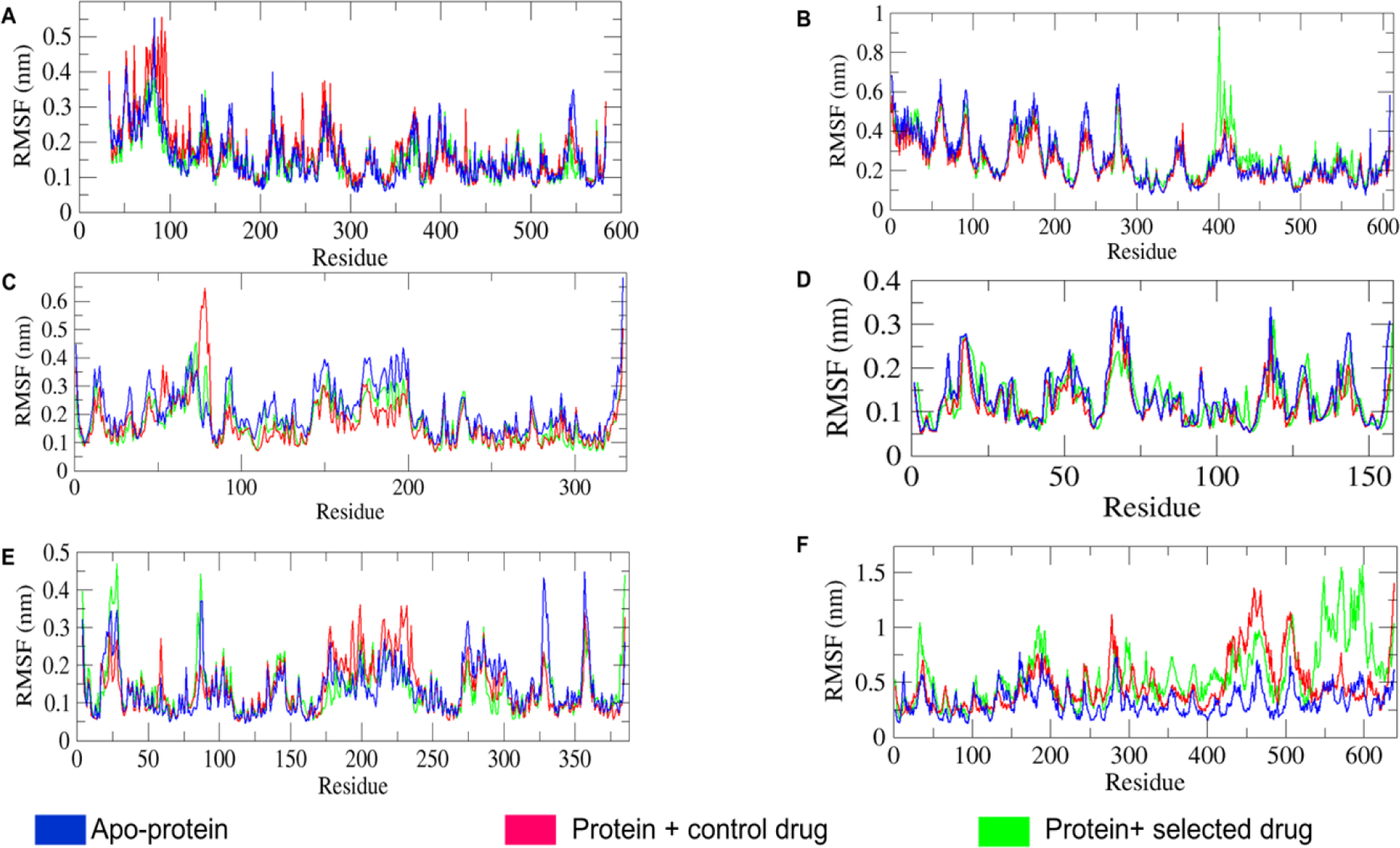
Comparative analysis of root-mean square fluctuation (RMSF) values of the consecutive apo-proteins and holo-proteins combinations (proteins with control drug and selected drug) mentioned in different subfigures (A) 1CX2 (B) 1MWU (C) 3N8D (D) 3SRW (E) 6KSI and (F) 3J25 combinations.

#### Structural compactness analysis

Structural compactness is analyzed by R_g_ (radius of gyration) values. Steady and less fluctuating R_g_ indicates a more compact protein structure. R_g_, the distance between the centre of mass of all protein atoms, is crucial for assessing stable protein folding [26]. As shown in Fig 7, owing to the volumetric size of proteins the R_g_ values for the apo-protein and holo-protein systems varied as follows: 2.44-2.54 nm for 1CX2, 3.55-3.80 nm for 1MWU, 2.10-2.40 nm for 3N8D, 1.54-1.65 nm for 3SRW, 2.05-2.18 nm for 6KSI, and 2.80-3.70 nm for 3J25. The comparative R_g_ trajectory of apo-protein and holo-protein systems in Figs 7A, 7B, and 7D shows that 1CX2, 1MWU, and 3SRW systems consistently maintained steady R_g_ values with moderate fluctuations. The holo-proteins closely overlapped with their corresponding apo-proteins, indicating that holo-proteins retained compactness similar to apo-proteins despite ligand interactions. Conversely, Figs 7C and 7E show that 3N8D-T-muurolol and 6KSI-T-muurolol exhibited greater compactness and integrity compared to their apo-protein and protein-control drug counterparts. The R_g_ trajectories for 3N8D-T-muurolol and 6KSI-T-muurolol were steadier and declined gradually after 40 and 60 ns, respectively, over the 100 ns MD run. Additionally, in Fig 7F, the 3J25 apo-protein showed steady behaviour with moderate R_g_ fluctuations compared to its holo-proteins. This indicates that 3J25 holo-proteins progressively lost compactness and stability in protein folding after successive ligand interactions.

**Fig 7.**
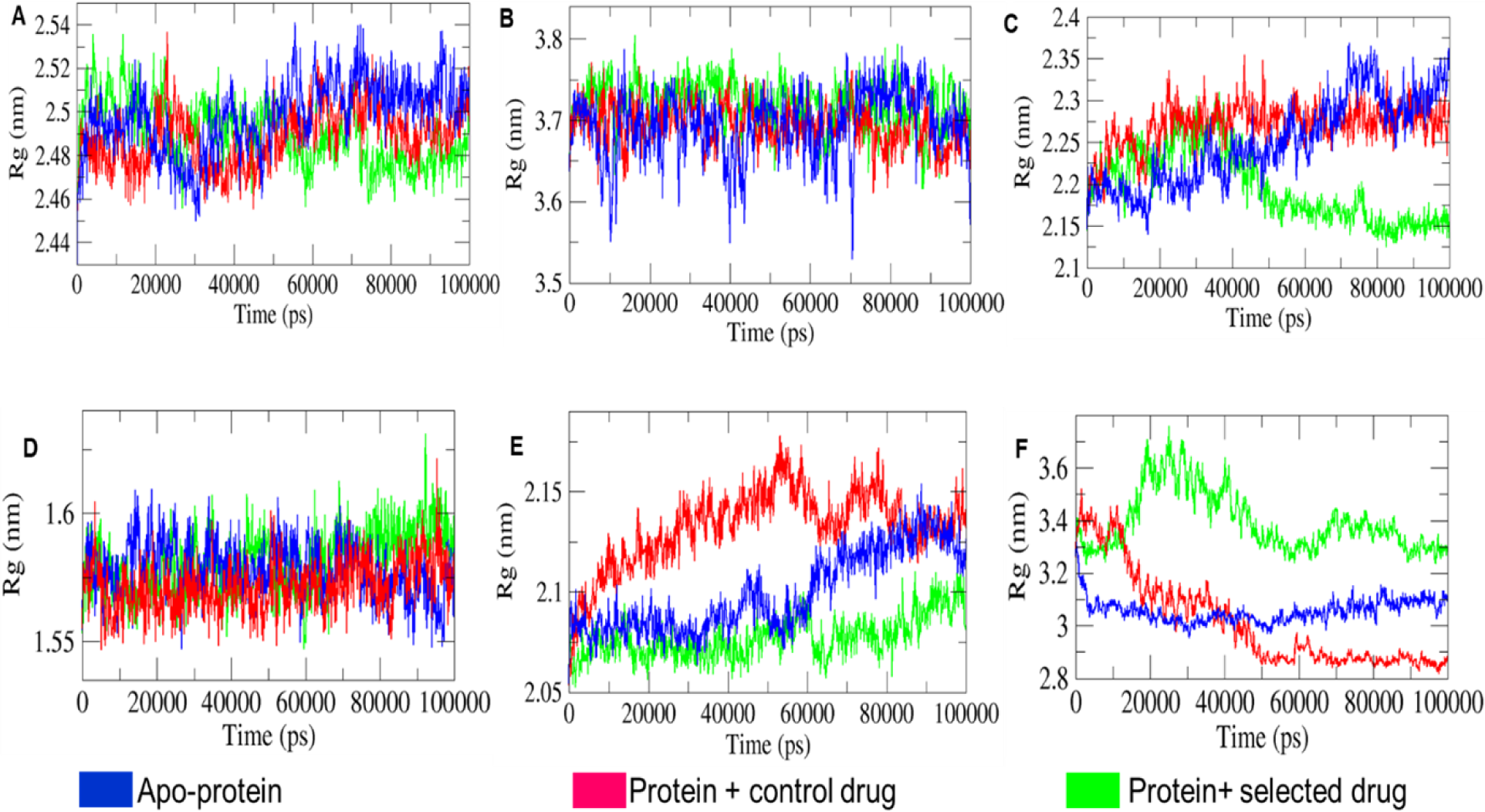
Comparative analysis of radius of gyration (R_g_) values of the respective combinations of apo-proteins and holo-proteins and the depicted combinations in different subfigures (A) 1CX2 (B) 1MWU (C) 3N8D (D) 3SRW (E) 6KSI and (F) 3J25 combinations.

#### SASA landscape analysis

SASA measures the protein surface area accessible to solvent molecules. With TIP3P water, hydrophilic apo-proteins showed increased SASA, while hydrophobic ones showed decreased SASA due to unfolding and folding. Ligand binding can shield surface residues, altering hydrophilicity and hydrophobicity, thus impacting the SASA and relative size of holo-proteins compared to apo-proteins [27]. As illustrated in Fig 8, SASA varied among apo-protein and holo-protein systems due to protein size: ∼(240-280) nm² for 1CX2, ∼(308-337) nm² for1MWU, ∼(160-185) nm² for 3N8D, ∼(90-100) nm² for 3SRW, ∼(165-195) nm² for 6KSI, and ∼(335-420) nm² for 3J25. In the 1CX2, 3N8D, and 6KSI systems, SASA of T-muurolol-bound holo-proteins significantly decreased during 50-100 ns, 40-100 ns, and 10-80 ns MD intervals, respectively, compared to their apo-proteins and protein-control drug complexes. This suggests substantial T-muurolol binding to surface residues during these intervals, leading to increased hydrophobicity, size shrinkage, and protein folding for 1CX2-T-muurolol, 3N8D-T-muurolol, and 6KSI-T-muurolol. Conversely, in Fig 8F, 3J25-T-muurolol showed increased SASA, indicating higher hydrophilicity than its apo-protein and protein-control drug complex during 10-85 ns. This suggests that ligand interactions led to greater hydrophilicity, protein unfolding, and relative size expansion for 3J25-T-muurolol compared to 3J25 apo-protein and 3J25-2-Oxazolidinone. Furthermore, substantial overlapping SASA trajectories and moderate alterations in time-dependent SASA values between apo-protein and holo-protein systems of 1MWU and 3SRW suggest comparatively less significant ligand-protein interactions in these systems. Except for 1MWU holo-proteins, the differing SASA profiles of T-muurolol-protein interaction complexes indicate that hydrophobic and van der Waals interactions predominantly contributed to binding affinity in T-muurolol holo-proteins, as supported by the MM-PBSA landscape of T-muurolol-protein interactions in Table 6.

**Fig 8.**
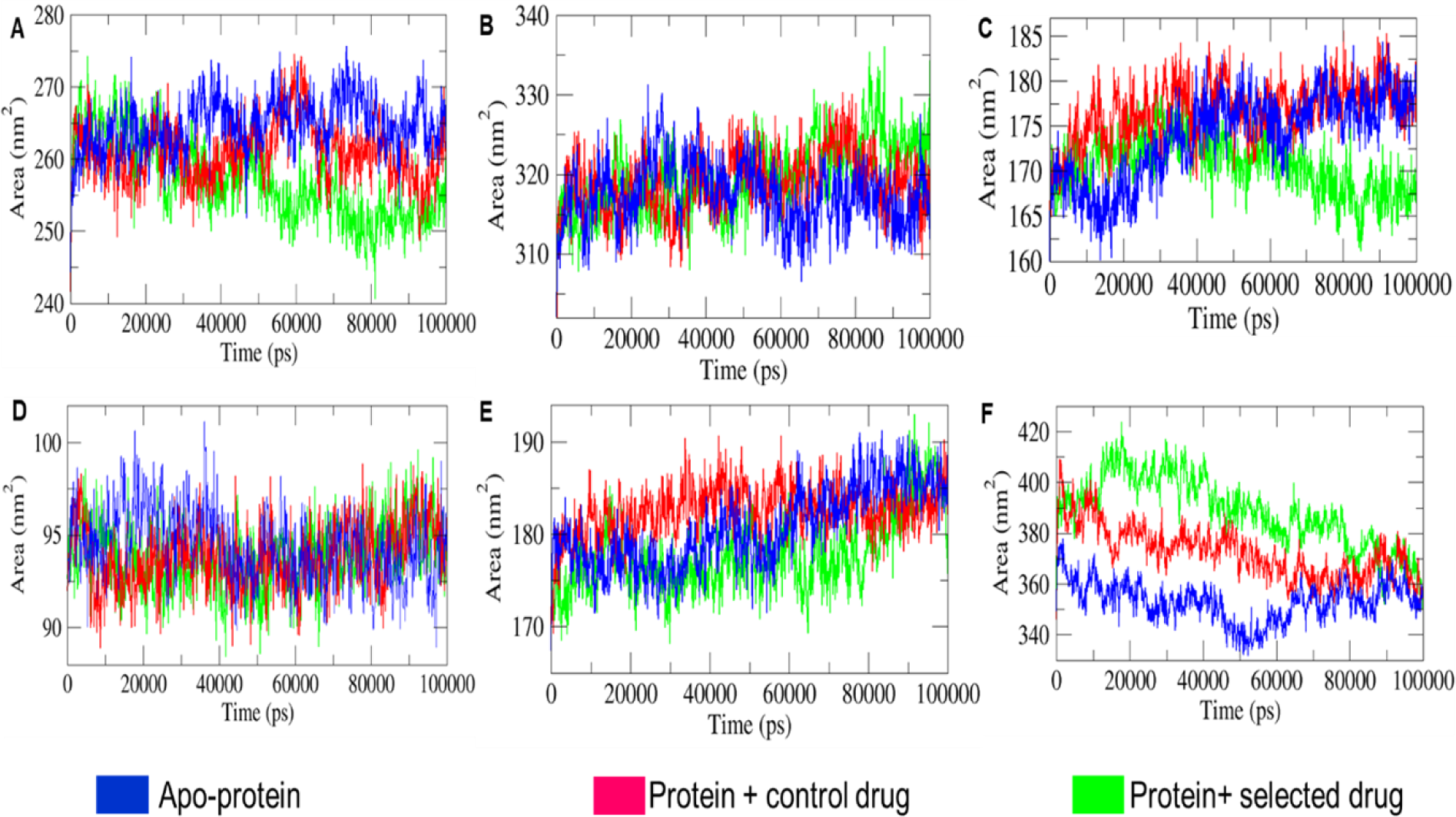
Comparative analysis of SASA (solvent accessible surface area) profiles of respective combinations of apo-proteins and holo-proteins and the depicted combinations in different subfigures (A) 1CX2 (B) 1MWU (C) 3N8D (D) 3SRW (E) 6KSI and (F) 3J25 combinations.

#### H bonds formation analysis

GROMACS analysis predicts ligand-protein complex stability by assessing hydrogen bond formation. Occupancy of H bonds in the interaction complexes of protein-ligands aids in heightening electrostatic energy flow [26]. In Fig 9, 6KSI-T-muurolol consistently formed one intermolecular hydrogen bond throughout the MD interval ∼(26-93) ns, indicating strong bond stability. Conversely, 1MWU-T-muurolol displayed intermittent bond formation at ∼85 ns, suggesting less stable interactions. Additionally, 1CX2-T-muurolol showed regular and irregular bond formation patterns at specific intervals, while 3N8D-T-muurolol exhibited intermittent bond formation across multiple MD time spans. 3SRW-T-muurolol maintained regular bond formation, whereas 3J25-T-muurolol formed bonds infrequently at various MD intervals.

**Fig 9.**
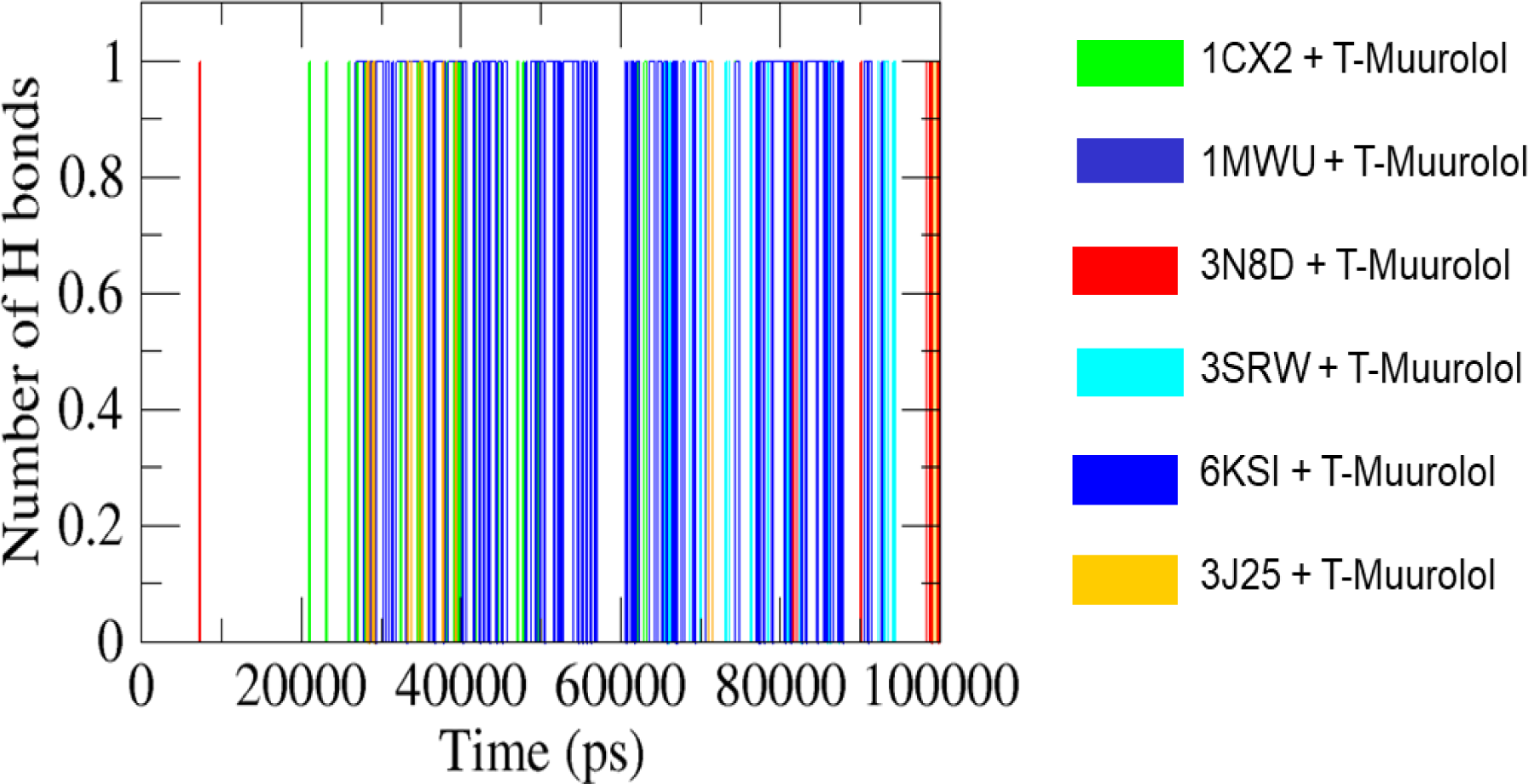
Illustration of the hydrogen bonds’ intensity up to MD time span of 100000 ps (picoseconds) formed between T-muurolol and respective proteins depicted in the figure itself.

#### Evaluation of MM-PBSA binding free energy decomposition

Table 6 presents the binding free energy decomposition for protein-ligand interactions involving T-muurolol with six macromolecules (1CX2, 1MWU, 3N8D, 3SRW, 6KSI, 3J25). The MM-PBSA binding free energies (ΔG*_BA_*) are approximately ∼(-39.48) KJ/mol (1CX2), ∼(-55.58) KJ/mol (1MWU), ∼(-21.39) KJ/mol (3N8D), ∼(-11.03) KJ/mol (3SRW), ∼(-50.64) KJ/mol (6KSI), and ∼(-5.01) KJ/mol (3J25). Van der Waals interactions contributed the most to ΔG*_BA_*, indicating that short-range dipole-dipole, permanent, and transient hydrophobic interactions are crucial in the protein-ligand binding stability. In addition to van der Waals energy, electrostatic and polar solvation energies reflect hydrogen bond formation, desolvation, and solvation effects among charged dipoles and polar and non-polar molecules in the MD environment. The protein-ligand complexes are surrounded by the polar solvent TIP3P (water) molecules and ions (K^+^ and Cl^-^). Thus, the conformational stability and folding of the proteins are heavily influenced by polar solvation and electrostatic energies. Polar solvation impacts the interactions by shielding the proteins’ polar hydrophilic residues, decreasing the hydrophobic binding affinity of ligands. This indicates that an increase in polar solvation energy negatively correlates with ligand-protein interactions, as it reduces the effectiveness of other binding energies (SASA, van der Waals, hydrophobic, and electrostatic). Consequently, higher polar solvation energy negatively impacts the overall stability of strong protein-ligand interactions [28]. Strong protein-ligand interactions influenced by polar solvation release positive energy, while SASA and electrostatic energies favour negative energy flow, indicating exothermic binding. Polar solvents like TIP3P enhance SASA interactions for hydrophilic surfaces, promoting protein folding and increasing hydrophobic interactions. Lower SASA in protein-ligand complexes correlates with better binding affinity, strengthening hydrophobic interactions and stabilising the complex [27, 29]. As shown in Table 6, the 1MWU-T-muurolol complex had the highest ΔG*_BA_* due to increased polar solvation energy and decreased electrostatic, van der Waals, and hydrophobic energies. This indicates significant polar solvation with TIP3P molecules interacting with surface hydrophilic residues, reducing SASA and other binding energies. Conversely, 6KSI maintained the strongest interaction with a ΔG*_BA_* of ∼(-50.64) KJ/mol through maximized hydrophobic and electrostatic interactions and minimized polar solvation effects over a 100 ns MD interval. This suggests that higher polar solvation energy and ΔG*_BA_* are not definitive indicators of strong protein-ligand interactions [28]. Thus, it is suggested that the combined contributions of van der Waals, electrostatic, SASA, and hydrophobic interactions in enhancing ΔG*_BA_* support robust ligand-protein interactions.

### Quantum chemical computational analysis

#### Frontier Molecular Orbital (FMO) analysis

Molecular Orbital (MO) analysis, especially the examination of Frontier Molecular Orbitals (FMOs) such as the Highest Occupied Molecular Orbital (HOMO) and the Lowest Unoccupied Molecular Orbital (LUMO), is essential for understanding a molecule’s electronic structure and reactivity. FMOs offer valuable insights into molecular interactions and help predict the chemical reactivity and stability of compounds. Fig 10A, display the HOMO and LUMO molecular orbitals for the optimized geometry of T-muurolol. The HOMO and LUMO energies for T-muurolol are -6.37 eV and 0.52 eV respectively with an energy gap (ΔE_Gap_) of 6.9 eV. In the FMO plots, the green and red lobes represent the orbital phases, with one colour indicating a positive phase and the other a negative phase.

**Fig 10.**
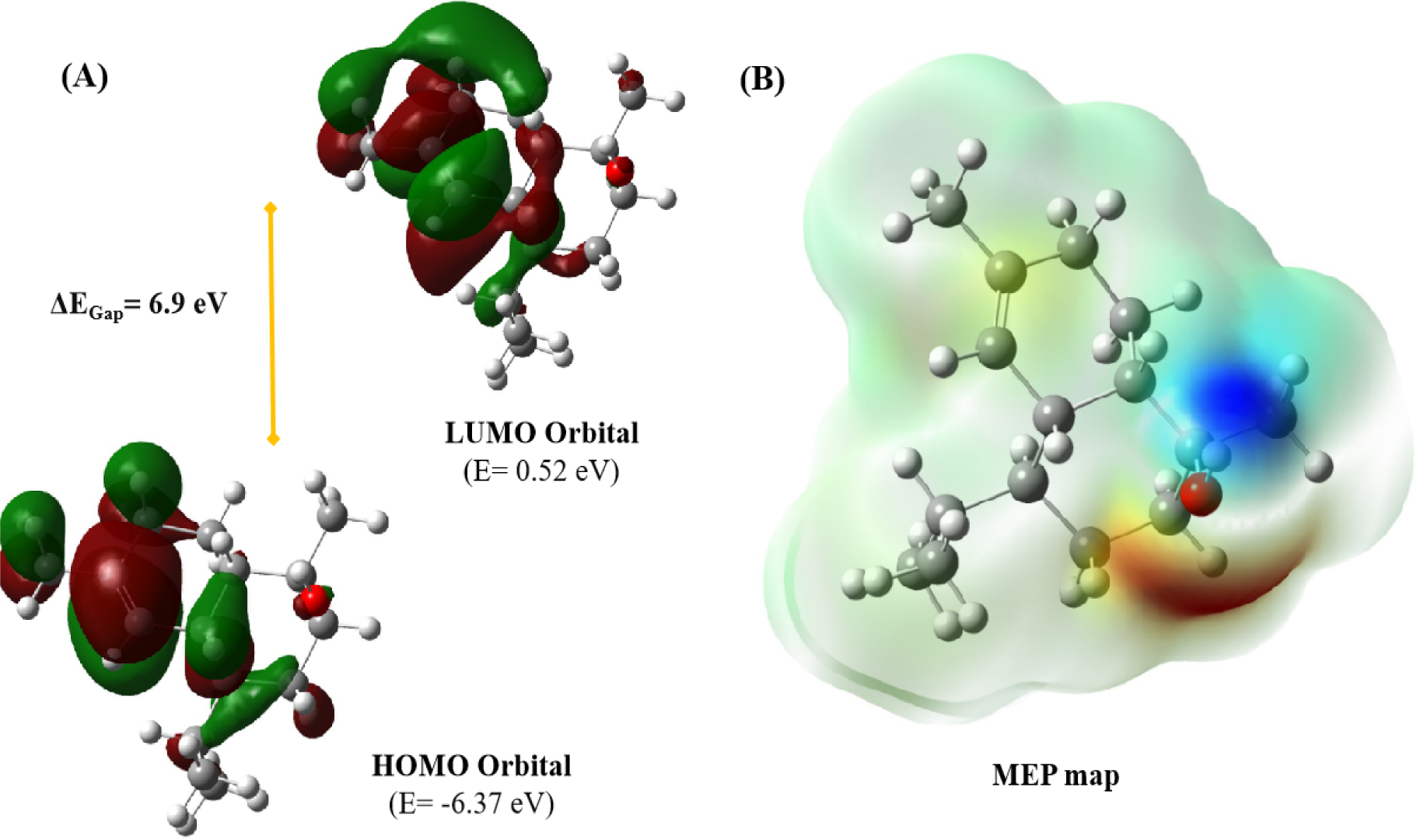
(A) HOMO and LUMO molecular orbitals of the T-muurolol molecule, along with their energy gap reported in electron volts (eV) using an iso-surface value of 0.02 electron. (B) Molecular Electrostatic Potential map of T-muurolol molecule (red atom: O; blue atom: N; grey atom: C; white atom: H). These plots are derived from optimizing the T-muurolol molecule using the DFT/B3LYP/6-311G(d,p) basis set level.

Hyperconjugation stabilizes molecular orbitals through overlap between an occupied orbital and a neighbouring electron-deficient orbital. As shown in Fig 10A, the HOMO is localized over the double bond region, typical of high electron density in alkenes, with some electron density around the hydroxyl group, indicating potential nucleophilic attack sites. The LUMO, on the other hand, shows electron density around the carbon atoms adjacent to the double bond and the hydroxyl group. The LUMO lobes, like those of the HOMO, also display phases (green and red) but are located in different regions of the molecule. The significant electron density around the double bond in both orbitals highlights this region as crucial for the molecule’s reactivity.

#### Molecular Electrostatic Potential analysis

A Molecular Electrostatic Potential (MEP) map visually represents the charge distribution across a molecule, highlighting positive and negative potential areas. This is useful for predicting molecular interactions, particularly electrostatic interactions. The MEP map of T-muurolol (Fig 10B) emphasizes the hydroxyl group as a key functional site due to its distinct polar regions. The map shows areas of positive potential (blue, 4.528 × 10-2-e) that are likely to attract nucleophiles and areas of negative potential (red, -4.528 × 10-2 e) that are likely to attract electrophiles, with the rest of the molecule being relatively neutral (green). The pronounced blue and red regions around the hydroxyl group indicate significant polarity, making this group in T-muurolol a reactive site for hydrogen bonding and polar interactions. The neutral green regions, covering most of the hydrocarbon skeleton, suggest that these parts of the molecule are less reactive but contribute to its overall shape and steric properties (Fig 10B).

#### Global chemical reactivity parameters

In computational chemistry, Koopmans’ theorem [30] serves as a powerful tool for estimating key electronic properties of molecules, enabling the derivation of essential global reactivity parameters. By applying this theorem, we can derive several crucial parameters that provide quantitative insights into the behaviour of the T-muurolol molecule, including electronegativity (◻ = -1/2 (E_HOMO_+E_LUMO_), chemical Potential (◻ = -◻), global Hardness index (◻ =1/2 (E_LUMO_-E_HOMO_)), global Softness index (σ=1/◻), global electrophilicity index (◻= ◻^2^/2◻). All parameters listed above provide (Table 7) a quantitative framework for predicting and rationalizing the chemical behaviour of T-muurolol molecules, aiding in the design and synthesis of new compounds with desired reactivity profiles.

**Table 7.** Molecular orbital energy and Koopman’s parameters for T-muurolol molecule obtained at DFT/B3LYP/6-311G(d,p) basis set level.

| List of parameters | Values calculated at the DFT level |
| --- | --- |
| Dipole moment (in debye or D) | 1.67 |
| $E_{\text{HOMO}}$ (in eV) | -6.37 |
| $E_{\text{LUMO}}$ (in eV) | 0.52 |
| electronegativity index ($\chi$, in eV) | 2.92 |
| chemical potential index ($\mu$, in eV) | -2.92 |
| global hardness index ($\eta$, in eV) | 3.45 |
| softness index ($\sigma$, in eV<sup>-1</sup>) | 0.29 |
| global electrophilicity index ($\omega$, in eV) | 1.24 |

## Discussion

T-muurolol is a bioactive sesquiterpenoid predominantly found as a major compound in the essential oils of several plants, including *Alpinia zerumbet*, *Pulicaria somalensis*, *Calocedrus formosana*, *Calendula officinalis*, and *Chamaecyparis obtusa* with reported antifungal, antibacterial, and antioxidant activities [12, 13,14, 31, 32]. With a documented history of bioactivity, T-muurolol could be an effective agent in inhibiting *S.aureus*-associated infections. For that, it was screened against various drug targets of *S. aureus* to evaluate its efficacy as an alternative antibacterial agent.

T-muurolol exhibited a strong binding interaction with *S. aureus* lipase (SAL) which is a triacylglycerol esterase, is a key virulence factor that degrades immune-responsive lipids, inhibits innate immune cell activation, and disrupts host immune recognition [33]. So, SAL is a major drug target for suppressing *S. aureus* pathogenesis. Strong interactions with SAL indicate its potency as an inhibitor. However, few SAL inhibitors have been introduced till now, suggesting that T-muurolol could be an effective agent for targeting SAL in further drug development. Similarly, dihydrofolate reductase, a key protein in the thymidine synthesis pathway essential for DNA synthesis, is crucial for bacterial survival [15]. The strong binding interaction of T-muurolol with this enzyme suggests its efficacy in inhibiting bacterial replication.

MDR *S. aureus* strains are major contributors to *S. aureus*-associated infections, posing significant management challenges and life-threatening risks to humans. T-muurolol has demonstrated strong molecular interactions with various drug targets specific to MDR *S. aureus* strains. For MRSA, T-muurolol targets penicillin-binding protein 2a, thereby inhibiting cell wall biosynthesis. It also exhibits a high binding affinity for D-Ala:D-Ala ligase, a critical drug target in VRSA. Additionally, T-muurolol interacts strongly with the RPP TetM protein in complex with the 70S ribosome, which is implicated in tetracycline resistance in TetRSA by mutating the interaction point of domain IV with the tetracycline binding site [34]. These interactions suggest that T-muurolol can inhibit TetRSA susceptibility by effectively targeting this protein. Furthermore, T-muurolol exhibits effective binding interactions with FAD-dependent NAD(P)H oxidase, which is involved in the ROS mechanism in the human body [35], and cyclooxygenase-2, which catalyzes the biosynthesis of pro-inflammatory prostanoids. These interactions establish its antioxidant and anti-inflammatory activities. These results consequently indicate the potential of T-muurolol as an effective agent for mitigating *S. aureus*-associated inflammation and cellular oxidative stress.

Moreover, MD simulation analysis elucidated the stability of T-muurolol interactions with selected proteins. RMSD analysis revealed enhanced backbone stability across all T-muurolol interactions compared to protein-control drug interactions. Similarly, RMSF analysis indicated stable interaction of T-muurolol with all proteins over the 100 ns MD timespan, surpassing control drugs. Except for the penicillin-binding protein 2a complex, differing SASA profiles of T-muurolol-protein interaction complexes suggested that hydrophobic and van der Waals interactions predominantly contributed to T-muurolol’s binding affinity. R_g_ values of proteins and protein-ligand complexes highlighted increased compactness and structural integrity in the presence of T-muurolol relative to control drugs. Hydrogen bonds remained consistent with docking scores, exhibiting minimal alterations in protein-ligand interaction dynamics. Furthermore, the recalculated binding energies of selected drugs using the MM-PBSA method after a 100 ns MD simulation demonstrated T-muurolol’s superior binding energy toward targeted proteins.

Additionally, quantum chemical structure analysis revealed a ΔE_Gap_ of 6.9 eV for T-muurolol, indicating greater stability and lower polarizability, resulting in higher chemical hardness [36]. A higher hardness value suggests lower reactivity [23], indicating the less reactive nature of T-muurolol. However, molecules with smaller HOMO–LUMO energy gaps are more interactive and inclined to participate in chemical events involving bond breaking or formation [37]. T-muurolol, with higher HOMO–LUMO values, signifies less anticipation in bond formations like covalent, hydrogen, and ionic bonds, correlating with molecular interaction studies showing fewer bond formations during protein-ligand interactions. A lower global electrophilicity index suggests a strong nucleophilic molecule, while a higher value indicates a strong electrophilic characteristic [38]. T-muurolol exhibits a moderate electrophilic nature with a value of 1.24 eV, indicating moderate facilitation of bond formation with biomolecules. Chemical potential reflects the electron escaping tendency from the molecule, with a lower value implying greater stability, indicating the greater structural stability of T-muurolol. Additionally, MEP analysis highlights the hydroxyl group in T-muurolol as the functional active site for hydrogen bonding and polar interactions.

Based on ADMET pharmacokinetics analysis, T-muurolol exhibits lower water solubility but possesses good absorption and distribution properties within the human body. Toxicity parameter analysis indicates that T-muurolol can be a promising drug candidate for inhibiting *S. aureus* bacterial-specific proteins, with very low toxicity observed. Succinctly, we propose that T-muurolol can be a promising antibacterial agent to combat *S. aureus* infections. While further in-vitro and in-vivo studies are necessary to confirm the outcomes and compare them with in-silico findings, this study lays the groundwork for subsequent investigations. These follow-up studies can provide insight into diverse prevention strategies for *S. aureus* infections.

### Conclusion

The present study explored the potential of T-muurolol as an alternative antibacterial agent against *S. aureus*. Several in-silico approaches such as molecular docking, MD simulation, quantum chemical structure analysis, and pharmacokinetic prediction have been utilized to screen the efficacy of T-muurolol as a drug. As a result, T-muurolol was identified as a potential antibacterial agent that can inhibit several *S. aureus-associated* bacterial proteins responsible for pathogenesis, bacterial survival, and multi-drug resistance. Along with it, T-muurolol is also identified as an antioxidant and anti-inflammatory agent that signifies its role in mitigating bacterial infection-associated inflammation and cellular oxidative stress in the human body. Further, quantum chemical structure analysis revealed its structural stability and less reactive nature. The results from the ADMET and bioactivity analysis strongly indicate that T-muurolol has the potential to act as a drug and demonstrate significant biological activities. Moreover, this study suggests that T-muurolol has the potential to counteract *S. aureus* infections as an alternative antibacterial solution. However, further detailed in-vitro and in-vivo experimental validation is required to confirm its activity.

## Materials and methods

### Target protein and ligand preparation

For molecular drug targets of *S. aureus*, several pathogeneses causing bacterial-specific proteins were identified such as V8 protease (PDB ID: 2O8L), signal Transduction Protein Trap (PDB ID: 4AE5), and *S. aureus* lipase (PDB ID: 6KSI). Along with them, several MDR *S. aureus*-specific proteins are also targeted such as beta-lactamase (PDB ID: 1GHI) and penicillin-binding protein 2a (PDB ID: 1MWU) for MRSA, Tet repressor protein (PDB ID: 2FJ1), RPP TetM in complex with 70S ribosome (PDB ID: 3J25), and QacA antiporter protein (PDB ID: 7Y58) for TetRSA, and D-Ala:D-Ala ligase (PDB ID: 3N8D) for VRSA respectively. Additionally, several universal bacterial proteins were identified, as described by Jianu et al. (2021) [15].

Further, to assess the bioactivity of the phytochemicals, we targeted several proteins involved in antioxidant and anti-inflammatory activities were targeted. For anti-inflammatory activities, cyclooxygenase-2 (PDB ID: 1CX2), lipoxygenase with protocatechuic acid (PDB ID: 1N8Q), hyaluronidase (PDB ID: 2PE4), lipoxygenase (PDB ID: 3V92), inducible nitric oxide synthase (PDB ID: 4CX7), and 11β-hydroxysteroid dehydrogenase 1 (PDB ID: 4YYZ) proteins were targeted whereas for antioxidant activities human cyclin-dependent kinase 2 complex (PDB ID: 1HCK), FAD-dependent NAD(P)H oxidase (PDB ID: 2CDU), glutathione peroxidase (PDB ID: 2F8A), and superoxide dismutase (PDB ID: 3HFF) were targeted.

The crystal structures of all proteins were retrieved from the RCSB PDB (accessed on March 12, 2024) and were prepared for docking studies by removing unwanted chains and residues using UCSF Chimera 1.16.

For ligand preparation, the 3D SDF structures of T-muurolol, ascorbic acid, ibuprofen, and 2-oxazolidinone were downloaded from PubChem (accessed on March 12, 2024). These compounds were imported into Avogadro 1.2.0 and subjected to energy minimization using the MMFF94 force field with the steepest descent algorithm. The structures were then converted to PDBQT format for docking.

### Molecular docking and binding analysis

AutoDock Tools 1.5.7 (Scripps Research Institute, La Jolla, CA, USA) was used to predict protein-ligand interactions. Water molecules were removed, and polar hydrogens and Kollman charges were added to the protein structures. The protein and ligand files were converted to PDBQT format. Active sites were selected based on blind-docking and enclosed within a 3D affinity grid centred on the active site residues. Docking was performed via command prompt as described by Gupta et al., 2023 [16]. Binding energies were recorded, and initial visualizations were done with PyMOL 2.5. Detailed interaction analysis was performed with LigPlot+ 2.2. Compounds with the best binding affinities were selected for further molecular dynamics simulation.

Furthermore, for cross-checking the binding affinities of the ligands with proteins, docking was performed in the CB-Dock2 online server (https://cadd.labshare.cn/cb-dock2/index.php, accessed on 12 March 2024).

### Pharmacokinetics prediction and bio-activity analysis

Pharmacokinetics parameters related to absorption, distribution, metabolism, excretion, and toxicity (ADMET) play a substantial role in the detection of novel drug candidates. To predict candidate molecules using in silico methods pkCSM (https://biosig.lab.uq.edu.au/pkcsm/prediction, accessed on 12 March 2024) online tool was used. Parameters such as AMES toxicity, maximum tolerated dose (human), hERG I and hERG II inhibitory effects, oral rat acute toxicities, hepatotoxicity, skin sensitization, and fathead minnow toxicity were explored. In addition to these, molecular weight, hydrogen bond acceptor, hydrogen bond donor, number of rotatable bonds, topological polar surface area, octanol/water partition coefficient, and number of violations of Lipinski’s rule of five were also surveyed by SwissADME (http://www.swissadme.ch/, accessed on 18 March 2024). The molecular properties and bioactivity score of the ligands were obtained from the molinspiration web server (https://www.molinspiration.com/cgi/properties, accessed on 18 March 2024).

### Molecular dynamics and simulation

The six best protein-ligand complexes from the molecular docking study, selected based on the lowest binding energy and optimal docked pose, were chosen for MD simulation. The macromolecular structures (PDB ID_1CX2, 1MWU, 6KSI and 3N8D) consisted of multiple subunits that were segregated and their additional homodomains had been removed via Discovery Studio Visualizer v20.1.0.19195 (https://www.3ds.com/products/biovia/discovery-studio) followed by the removal of adjacent heteroatoms via similar fashion. Then the modified structures were further modified to become compatible with in-silico virtual screening and molecular dynamic simulation (MDS). A comparison map of the dynamic characteristics of targated proteins and their protein-ligand complexes using GROMACS 2020.1 according to described methods by Kandasamy et al. (2022) [17]. In this study, The PDB structures of the proteins were transformed to gmx (gromacs) format using the CHARMM36 force field (http://www.charmm-gui.org/). The parameter files for the docked complexes were created using the guidelines outlined in the GROMACS course. The topology and parameter files for protein and ligand were generated on the CHARMM-GUI server, and selected protein-ligand complex files were prepared using the CHARMM-GUI Multicomponent Assembler (MCA). The cube enclosing the system was sized based on ’Calculated Volume Fraction,’ ’Minimum Recommended Size Length,’ and ’Maximum Volume Fraction,’ with a 20 Å buffer to prevent periodic image overlap during MD simulation. Following the final system size determination, the solvent composition was calculated, and the system was solvated with TIP3P water and 150 mM K^+^ and Cl^-^ ions per the Solution Builder protocol. GROMACS files for proteins and protein-ligand complexes were extracted from CHRMM-GUI files for 100 ns (nanoseconds) with 2 femtoseconds (fs) steps.

The analysis of root means square deviation (RMSD), root means square fluctuations (RMSF) of backbone atoms, solvent accessible surface area (SASA), radius of gyration (R_g_) and hydrogen bond (HB) formation were analyzed after the successful completion of MD simulation of specific protein-ligand interaction complexes with a time interval of 20 ns throughout the whole 100 ns of MD trajectory. All the graphs (RMSD, RMSF, SASA, R_g_, HB,) were plotted by using Qtgrace (https://qtgrace.sourceforge.io/).

### Binding free energy analysis

Apart from in-silico docking-based binding affinity analysis between proteins and specific ligands, the strength of protein-ligand interactions was further assessed using MD simulation-specific binding free energy analysis. This was accomplished by calculating the binding free energies with the MM-PBSA (Molecular Mechanics Poisson-Boltzmann Surface Area) method. Using this method, the GROMACS function “g_mmpbsa” captured MD trajectory files of protein-ligand interaction complexes throughout 100 ns at 5 ns intervals by using the following equation ΔG*_BA_* = ΔE*_MM_*+ ΔG*_PBSA_* – TΔS*_MM_* Where ΔG*_BA_*, ΔE*_MM_*, ΔG*_PBSA_*, and TΔS*_MM_* denote as average free energy, average molecular mechanics energy, solvation energy, and solute configuration entropy respectively [17].

### Computational chemical structure analysis

The chemical structure of the T-muurolol molecule was optimized using the DFT/B3LYP method with the 6-311G(d,p) basis set in the Gaussian-09 program (Gaussian 09, Revision A.02)[18]. This optimized geometry was then used to generate the Frontier Molecular Orbitals (FMOs) and Molecular Electrostatic Potential (MEP) plots for the T-muurolol molecule. Additionally, the chemical potential of the structure was examined using DFT.

## Acknowledgements

The authors acknowledge the project “e-Infrastruktura CZ” (e-INFRA ID:90254) provided within the program Projects of Large Research, Development and Innovations Infrastructures in the Czech Republic for suppling computational resources.

